# Optimal colors can predict luminosity thresholds in natural scenes

**DOI:** 10.1101/2025.08.07.669217

**Authors:** Killian Duay, Takehiro Nagai

## Abstract

Luminosity thresholds define the luminance boundary at which a surface color shifts in appearance from being perceived as an illuminated surface to appearing self-luminous. Previous research suggests that the human visual system infers these thresholds based on internal references of physically realizable surface colors under a given illumination, referred to as the physical gamut. A surface is perceived as self-luminous when its luminance exceeds the upper limit of this empirically internalized gamut. However, the precise structure and boundaries of these gamuts remain uncertain. Optimal colors, which represent theoretical surface reflectances under specific illuminants, have been shown to provide an effective model for visualizing and computing the physical gamuts. In prior studies, optimal colors have successfully predicted luminosity thresholds; however, these findings were limited to highly simplified, abstract stimuli. Whether this framework generalizes to more naturalistic viewing conditions has remained an open question. In the present study, we demonstrate that the theory of an internal reference in the form of an empirically constructed physical gamut, visualized through optimal colors, remains valid under more natural conditions. Our results confirm that optimal colors can still accurately predict luminosity thresholds in such settings. Moreover, our findings suggest that the luminosity thresholds encompass both self-luminosity and naturalness concepts. Subsequently, this may imply that the notion of physical gamut could envelope both concepts as well and could be defined as “all physically possible colors in a scene for an object that does not emit light.” These insights can have profound potential implications for both applied fields (i.e., XR or projection mapping) and fundamental science (e.g., understanding human visual processing mechanisms).

## Introduction

Visual objects can be perceived in two fundamentally different ways: either as illuminated surfaces reflecting light or as self-luminous sources emitting light. These perceptual distinctions are known as modes of appearance [1]. Despite the fact that the light entering the eye is similar in both cases, observers typically have no difficulty distinguishing between an object that is lit and one that appears to emit light [2]. Yet, when an object’s appearance lies near the boundary between these two modes, perception can become ambiguous. This boundary is referred to in the literature as the luminosity threshold: the point at which an object transitions from being perceived as illuminated to being seen as self-luminous. Understanding this threshold has been the focus of numerous studies aiming to pinpoint where this perceptual shift occurs, what factors influence it, and how the visual system processes such information to form coherent interpretations of object appearance.

Classical studies have investigated various factors influencing the luminosity thresholds, particularly in the context of chromatic and achromatic stimuli. Evans and Swenholt [3–6] demonstrated that, for chromatic stimuli, the luminosity threshold is negatively correlated with stimulus purity: the higher the purity, the lower the luminance required for a surface to appear fluorescent. In this context, fluorescence was defined as a color appearance lacking a gray component and perceived as self-luminous. Later, Ullman [7] attempted to account for luminosity thresholds using scene-based heuristics and image statistics such as maximum scene intensity, absolute stimulus intensity, intensity comparison with the average intensity of the scene, and local or global contrast, but concluded that none of these factors alone could reliably predict the thresholds. For achromatic surfaces, Bonato and Gilchrist [8] found that a stimulus begins to appear luminous when its luminance reaches approximately 1.7 times that of a surface perceived as white under the same illumination, regardless of whether an actual white reference is present in the scene. Expanding the theoretical framework, Speigle and Brainard [9] hypothesized that the perception of luminosity arises when the visual system infers a surface reflectance spectrum that lies outside the gamut of physically realizable surfaces under a given perceived illuminant. According to their model, a patch is perceived as self-luminous when its color is incompatible with any surface color that could exist under the estimated illumination conditions. Their work verified this hypothesis and demonstrated that the threshold for perceiving luminosity depends not only on the chromaticity of the stimulus, but also on the chromaticity of the illumination. Notably, this was the first introduction of the concept of a gamut of physically realizable surfaces under a specific illumination. For simplicity and conciseness, we refer to this concept as the *physical gamuts* in this study. Finally, Uchikawa et al. [10] showed that, for colored surfaces, perceived brightness, rather than luminance, plays a critical role in determining the transition between surface-color and aperture-color (self-luminous) modes. They also reaffirmed the negative relationship between stimulus purity and luminosity threshold. While these foundational studies have significantly advanced our understanding of the factors influencing luminosity perception, they fall short of offering a complete account of the underlying mechanisms.

More recently, Morimoto et al. [2] revisited the concept of the physical gamut (the range of colors physically realizable under a given illumination) by further advancing the idea that judgments of self-luminosity are based on an internal reference encoded by the human visual system. Unlike Speigle and Brainard [9], who derived the physical gamuts from empirical measurements using the reflectance spectra of real-world Munsell patches, Morimoto et al. [2] proposed a more theoretical framework grounded in the concept of optimal colors [11, 12]. Optimal colors represent, for each chromaticity, the highest possible luminance achievable through reflected light under a specific illuminant. Though theoretical, these optimal colors define the upper luminance boundary of surface colors and serve as an effective model for estimating the limits of what can be encountered in natural scenes. According to Morimoto et al. [2], the construction of the physical gamut as an internal reference in the human brain is shaped through empirical observations of the world and the surface colors encountered in daily life. Since optimal colors represent the highest luminance that can be produced and observed for any given chromaticity and surface color, this theoretical model would accurately represent and predict the boundaries of what can be observed in the real world and, consequently, the internal reference built by the visual system. They further propose that the human visual system uses this internal reference when judging whether a surface appears self-luminous or illuminated. This idea builds on previous work in color constancy [13–16], in which similar models of internalized optimal colors distributions successfully accounted for observers’ illuminant estimations across a range of viewing conditions. Applying this framework of optimal colors to the domain of luminosity perception, they hypothesized that the visual system determines whether a surface appears self-luminous by comparing its appearance to the internalized physical gamut: if the color lies within the gamut, it is perceived as an illuminated surface; if it lies outside, it is perceived as self-luminous. This hypothesis was tested and supported through three experiments. The results showed that luminosity thresholds peaked around the chromaticity of the test illuminants and decreased with increasing chromatic purity. Importantly, the loci of these thresholds closely matched the boundaries predicted by the physical gamut derived from optimal colors theory, and remained nearly invariant to changes in the surrounding color distribution. These findings suggest that while contextual cues may be essential for estimating the illuminant, they play a limited role in the final luminosity judgment once the illuminant is inferred. Although the notion of an internal physical gamut was introduced by Speigle and Brainard in a practical approach [9], Morimoto et al. [2] demonstrated that optimal colors currently offer the most precise and predictive model for the computation and visualization of the physical gamut internalized within the visual system.

Despite the valuable insights offered by Morimoto et al. [2], their study presents certain limitations that restrict the generalizability of its conclusions to real-world perception. Specifically, the experiments relied on highly simplified stimuli: 2-degree circular colored test fields embedded within arrays of overlapping colored circles. These abstract stimuli bear little resemblance to the complex visual environments encountered in daily life, making it difficult to extrapolate the findings to naturalistic scenarios. As a result, it remains unclear whether the mechanisms proposed in their optimal colors model are indeed employed by the human visual system when judging the appearance of objects and surfaces in realistic contexts. Prior research suggests that naturalistic features such as shading and surface texture significantly influence the perception of self-luminosity. For example, a uniformly colored circle may appear self-luminous when isolated, but the same stimulus can lose its luminous appearance when presented as a shaded 3D sphere, and even more so when realistic surface textures are added [17]. More broadly, shading alone has been shown to reduce perceived luminosity [18], and even minor surface irregularities, such as scratches, can disrupt the impression of self-luminosity and lead observers to interpret the object as a standard reflective surface [17, 18]. To further understand the processes underlying luminosity judgments and to robustly validate the optimal colors theory, it is essential to test whether the model proposed by Morimoto et al. [2] remains valid under more natural conditions. Such investigations are not only important for advancing theoretical understanding but also hold potential practical applications in areas like extended reality (XR) and projection mapping, where perceived self-luminosity plays a crucial role in visual realism.

In this study, we aimed to evaluate the physical gamut theory visualized by optimal colors developed and tested by Morimoto et al. [2], but using more naturalistic visual stimuli. For this purpose, we closely replicated the original methodology and experimental design employed in their study, ensuring consistency with their approach. The only modifications we intended to make concerned the stimulus, which we sought to gradually evolve toward a more naturalistic and realistic form. By doing so, this design would allow us to assess whether the optimal colors can also predict luminosity thresholds in natural stimuli, as it successfully did in more abstract ones. To this end, we designed and conducted a series of three psychophysical experiments wherein observers adjusted the luminance of a target within a scene until it appeared self-luminous; these settings were then compared against predictions derived from the optimal colors model. The three experiments utilized natural scene background images but differed in the type of the target to adjust, increasing in naturalness across experiments: (1) a 2D uniform circle superimposed on the background in Experiment 1, a shaded matte 3D sphere superimposed on the background in Experiment 2, and (3) an “in-picture” target in Experiment 3, where the pixels of an object within the original background image were directly manipulated. Based on aforementioned prior research, we hypothesized that the theory of the optimal colors representing physical gamuts and predicting luminosity thresholds would be: (1) confirmed in Experiment 1, as it has already been demonstrated that the background does not influence the results as long as it allows for an accurate estimation of the chromaticity and intensity of the illuminant [2]; (2) confirmed in Experiment 2, albeit with relatively higher observers’ settings than in Experiment 1 to compensate for the perceived reduction in luminosity induced by the shading of the sphere; and (3) refuted in Experiment 3, as the in-picture targets consist of natural objects with textures that disrupt the perception of self-luminosity. Remarkably and against all expectations, our results demonstrated that the theory remained valid to predict luminosity thresholds across all three experiments, successfully predicting and explaining luminosity thresholds in even the most naturalistic conditions. As anticipated, many observers reported that they could not directly perceive the targets as self-luminous in Experiments 2 and 3 but relied on an alternative strategy to assess luminosity thresholds: they judged them based on “unnatural brightness”. They reported that they were able to determine when the luminosity is excessively too high to be plausible or realistic, based on an impression of naturalness. Although they shifted their judgment criterion in order to be able to perform the task, their results were completely identical to those of observers who could directly judge self-luminosity. Moreover, the match with the optimal colors theory predictions was observed equally well in both cases. These results further support the theory of an internal reference in the form of a physical gamut, visualized through optimal colors, which was proposed by Morimoto et al. [2] and validated here under more natural conditions. Moreover, these findings suggest that luminosity thresholds may integrate both the concepts of self-luminosity and naturalness. Consequently, this may imply that the notion of the physical gamut similarly encompasses these two dimensions and could be defined as “all physically possible colors in a scene for an object that does not emit light.” Such a gamut may serve as an internal reference against which we evaluate whether a color appears plausible as a surface property; if it falls outside this range, it may be perceived as self-luminous or as unnatural. As stated before, these findings can have profound potential implications for both applied fields (e.g., XR or projection mapping) and fundamental science (i.e., understanding human visual processing mechanisms).

Before proceeding further, it is important to clarify the definitions of key concepts and the relationships between them. This terminology, drawn from the literature cited above, can be somewhat counterintuitive, and we aim to eliminate any potential ambiguity. The term luminosity threshold refers to the perceptual boundary between two modes of appearance: surface color, in which an object is perceived as illuminated and reflecting light (e.g., a painted wall), and self-luminosity, in which an object is perceived as emitting its own light (e.g., a light bulb). To explain where this boundary lies, Speigle and Brainard [9] introduced the concept of the physical gamut: the set of colors corresponding to physically realizable surface reflectances under a given illumination. The idea is straightforward: a given surface appears self-luminous when its luminance exceeds the upper-limit luminance of the physical gamut internalized in the visual system. The internalization of the physical gamut in the human brain occurs through empirical observation of the real world and the surface colors we encounter in everyday life. However, since we do not have access to an exhaustive database of all surface reflectances encountered in daily life, the exact structure and boundaries of these internal gamuts remain uncertain. This concept was later expanded upon by Morimoto et al. [2], who demonstrated that this gamut can be visualized by the theory of optimal colors: a theoretical construct representing the highest luminance attainable by a surface color for each chromaticity under a specific illuminant. The optimal colors model provides a computational means of estimating the physical gamut and visualizing how it may be internalized in the brain. Among existing frameworks, the optimal colors theory stands as the most recent and effective approach for supporting the concept of physical gamuts and explaining the perceptual judgments associated with luminosity thresholds. However, Morimoto et al. [2] demonstrated this theory using only abstract stimuli. In the present study, we provide evidence that the theory also holds for more naturalistic stimuli.

## General methods

### Observers

The observers were two females and six males. All were undergraduate or graduate students from Institute of Science Tokyo. All observers passed the Ishihara color vision test and had normal or corrected-to-normal visual acuity. All experiments were designed in accordance with the Declaration of Helsinki and was approved by the Ethical Review Committee of Institute of Science Tokyo. Informed written consent was obtained from all observers after explaining the details of experimental protocols.

### Apparatus

The experiments took place in a darkroom. Stimuli were displayed on an OLED monitor (PVM-A250, Sony, Japan) with a 10-bit per channel color resolution, a spatial resolution of 1920 × 1080 pixels, a 25-inch screen diagonal, and a 60 Hz refresh rate. To ensure accurate luminance and chromaticity presentation, the display was carefully calibrated using a spectroradiometer (Specbos1211-2, JETI Technische Instrumente GmbH, Germany) and a colorimeter (ColorCAL II, Cambridge Research Systems, UK). The monitor was connected to a laptop (MacBook Air, Apple, USA; Apple M2, macOS Sequoia 15.2), and the experiments were controlled by a custom program written in PsychoPy 2024.2.4. Observers’ heads were stabilized using a chinrest at a 40 cm viewing distance, and responses were collected via a numeric keypad and a mouse.

### Color computation

In color science, the CIE 1931 color space provides a standardized method to quantify color based on human vision. To compute colors in this space, three key components are required [19]: the spectral reflectance of a surface, the spectral power distribution (SPD) of the illuminant, and the CIE 1931 color matching functions (CMF), which represent the average human eye’s sensitivity to different wavelengths (*2° Standard Observer* in the present study). The spectral reflectance describes how much light a surface reflects at each wavelength, while the illuminant’s SPD defines the intensity of incident light across the spectrum. The reflected light is computed by multiplying the reflectance with the illuminant at each wavelength. This product is then integrated across the visible spectrum, weighted by the CMFs, to yield the final color expressed in tristimulus values *(X, Y, Z)*, from which the chromaticity coordinates *(x,y)* and the luminance *Y* can be derived. Hyperspectral images contain detailed spectral reflectance information for each pixel, enabling accurate per-pixel color computation under any defined illuminant. For more details, see previous literature [19, 20]. This is the method we used to generate the colors and stimuli in this study.

### General stimulus

The stimulus was presented at the center of a completely black background of 0 candelas per square meter (cd/m^2^), made possible by the OLED display used in our setup, as illustrated in Fig 1E. The stimulus subtended approximately 13.26° horizontally and 10.00° vertically, and was composed of two main elements: a background and a target.

**Fig 1.**
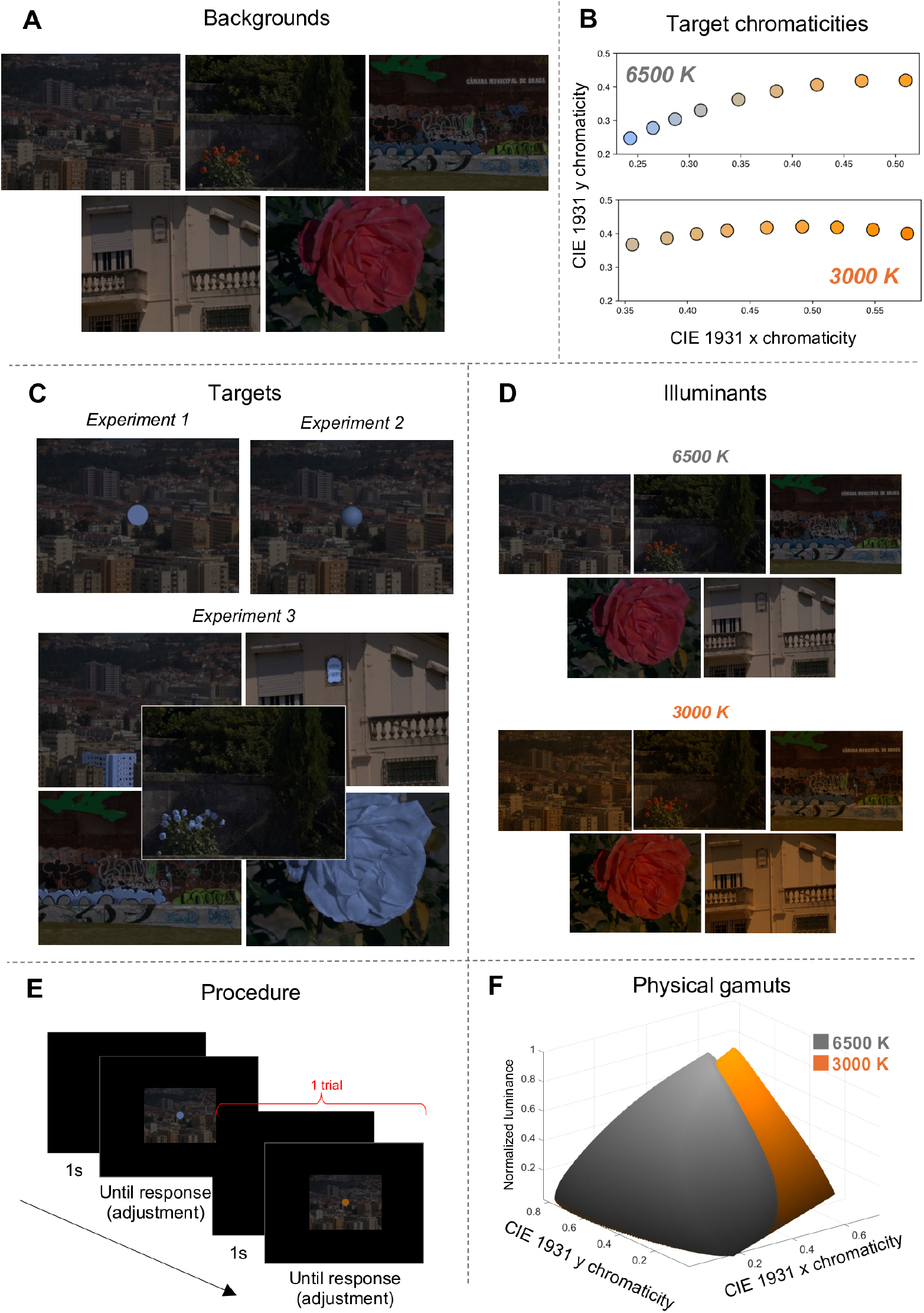
Methods of the three experiments. **A**: backgrounds of the experiments. **B**: test chromaticities of the targets. **C**: targets of Experiments 1,2, and 3. **D**: illuminants of the experiments. **E**: procedure of the experiments with an example of two trials. **F**: normalized physical gamuts of illuminants 6500 Kelvin (K) and 3000 K.

Five different backgrounds were used (see Fig 1A), each derived from hyperspectral images obtained from a publicly available dataset [21]. These images were selected to span a variety of natural scenes, including urban environments, vegetation-rich areas, and mixed-content settings, thereby sampling diverse distributions of natural colors. To simulate different lighting conditions, we applied two illuminants corresponding to distinct correlated color temperatures (CCTs): a neutral daylight-like illuminant of 6500 Kelvin (K) and a warmer, orangish illuminant of 3000 K. The backgrounds were generated by multiplying the hyperspectral data with the spectral power distributions of Planckian radiators at 6500 K and 3000 K, respectively, as shown in Fig 1D. This results in five available backgrounds for each of the two illuminant conditions used in our experiments: 6500 K and 3000 K. The illuminant intensities were scaled such that a perfect white surface (reflecting 100% of all wavelengths) under either lighting condition would yield a luminance of 15 cd/m^2^. Importantly, the same set of backgrounds was used across the three experiments.

The targets differed between the three experiments, as shown in Fig 1C. The target was a 2D circle in Experiment 1, a 3D sphere in Experiment 2, and an object embedded within the image in Experiment 3. The targets for each experiment are described in more detail in the respective sections below.

The target colors were the same for all three experiments. They were defined by a chromaticity and luminance values in CIE 1931 color space. Across all conditions, the target could assume one of nine distinct chromaticities per trial, as illustrated in Fig 1B. Only one chromaticity was applied at a time, and it was uniformly mapped onto the target (as exemplified by the blue chromaticity in Fig 1C; the chromaticity coordinates (*x,y*) of these pixels were transformed into the test chromaticities). In a trial, the same chromaticity was applied uniformly to the target and remained fixed (it did not vary with the observers’ adjustments). This was the case for all three experiments. With the intention of replicating the experimental conditions of Morimoto et al. [2], we followed the same strategy as them to select natural colors: the chromaticities were selected to lie along the black body locus and corresponded to Planckian radiator CCTs of 2000 K, 2400 K, 2900 K, 3500 K, 4300 K, 5600 K, 7200 K, 10000 K, and 20000 K. These were chosen to be approximately equally spaced along the locus on the CIE 1931 chromaticity diagram. The spectral power distributions of these nine Planckian radiators were multiplied with the spectral distributions of the two illuminants conditions used in the experiments (3000 K and 6500 K), yielding the final test chromaticities. Notably, the chromaticity resulting from the 5600 K Planckian radiator closely matched the white point of both experimental illuminants.

The target luminance was directly set and adjusted by the observers’ during each trial task. Since the targets differed across experiments, the method for applying the observer’s selected luminance value to the target also varied across the three experiments. The exact methods used to manipulate luminance in each experiment are presented in the respective sections below.

All resulting stimuli were generated in the CIE 1931 color space and then converted to corresponding RGB values for the display used in our experiment. Final color rendering on the monitor was ensured through a color calibration procedure, based on prior spectral measurements and display characterization.

### General procedure

Fig 1E illustrates two example trials from the experimental procedure. Each trial began with a 1-second blank and black screen, followed by the presentation of the stimulus, which remained visible until the observer judged their task to be complete and validated their response using the numeric keypad. The task consisted of adjusting the luminance of the target using a combination of mouse controls. Specifically, the mouse scroll wheel allowed for luminance changes in steps of 1.0 cd/m^2^, while the left and right mouse buttons provided finer adjustments, decreasing or increasing luminance in 0.5 cd/m^2^ increments, respectively. The observer’s goal was to determine and set the luminance at which the target began to appear self-luminous.

During a trial, only the luminance of the target was modified; the chromaticity remained fixed. Technically, this was achieved by converting the RGB representation of the stimulus back into the CIE 1931 color space, adjusting the luminance component, and converting it back into RGB for display. This process was performed in real-time and was imperceptible to the observer, ensuring a seamless visual experience.

Each session consisted of 45 trials, corresponding to all combinations of five background scenes and nine target chromaticities. The order of the five backgrounds was randomized, and within each background block, the chromaticities were presented in random order. Importantly, each session was associated with a single illuminant condition and did not mix stimuli from different illuminant sets. For example, a session dedicated to the illuminant 6500 K featured only the five backgrounds and nine chromaticities computed under that specific illuminant. The exact details of the sessions differed slightly across experiments and are outlined in the respective sections below.

Each session began with a 2-minute dark adaptation phase during which the screen remained entirely black, followed by a 30-second adaptation phase to the experimental illuminant and background content. During this adaptation period, the five background images used in the current session were displayed in a continuous loop, each shown for two seconds.

### Optimal colors and physical gamut

Optimal colors are defined as surfaces whose spectral reflectance functions consist exclusively of 0% and 100% reflectance values across the wavelength spectrum, with at most two abrupt transitions between these extremes [2, 13, 16]. In the real world, however, no material can achieve 100% reflectance at any wavelength due to physical limitations. Consequently, optimal colors exhibit higher luminance than any real surface possessing the same chromaticity, and thus, no naturally occurring surface can exceed the luminance of these theoretical distributions at a given chromaticity. Therefore, by computing all possible optimal reflectance functions and multiplying them spectrally with a given illuminant, one obtains the complete set of optimal colors that can be produced under that given lighting. This set forms the physical gamut for the specific illuminant. In our study, we generated optimal colors in the CIE 1931 color space following established methodologies [2, 13, 16]. Specifically, we produced approximately 90,000 optimal spectral reflectance functions, corresponding to every possible combination of reflectance transitions that satisfy the criteria of optimal colors. Each spectrum was then multiplied with the spectral power distributions of the Planckian radiators at 6500 K and 3000 K. The resulting sets of optimal colors, forming the physical gamuts under each illuminant condition used in our study, are shown in Fig 1F. The specific optimal colors corresponding to the chromaticities of the experimental targets, used for plotting and analyzing the 2D loci in Fig 2, were selected from within this set. It is important to note that the physical gamuts displayed in Fig 1F are expressed in normalized luminance values. To obtain absolute luminance, these values must be multiplied by the scene’s illuminant intensity (the intensity used to compute the spectral product from the hyperspectral images), which in our experiments was fixed at 15 cd/m^2^.

**Fig 2.**
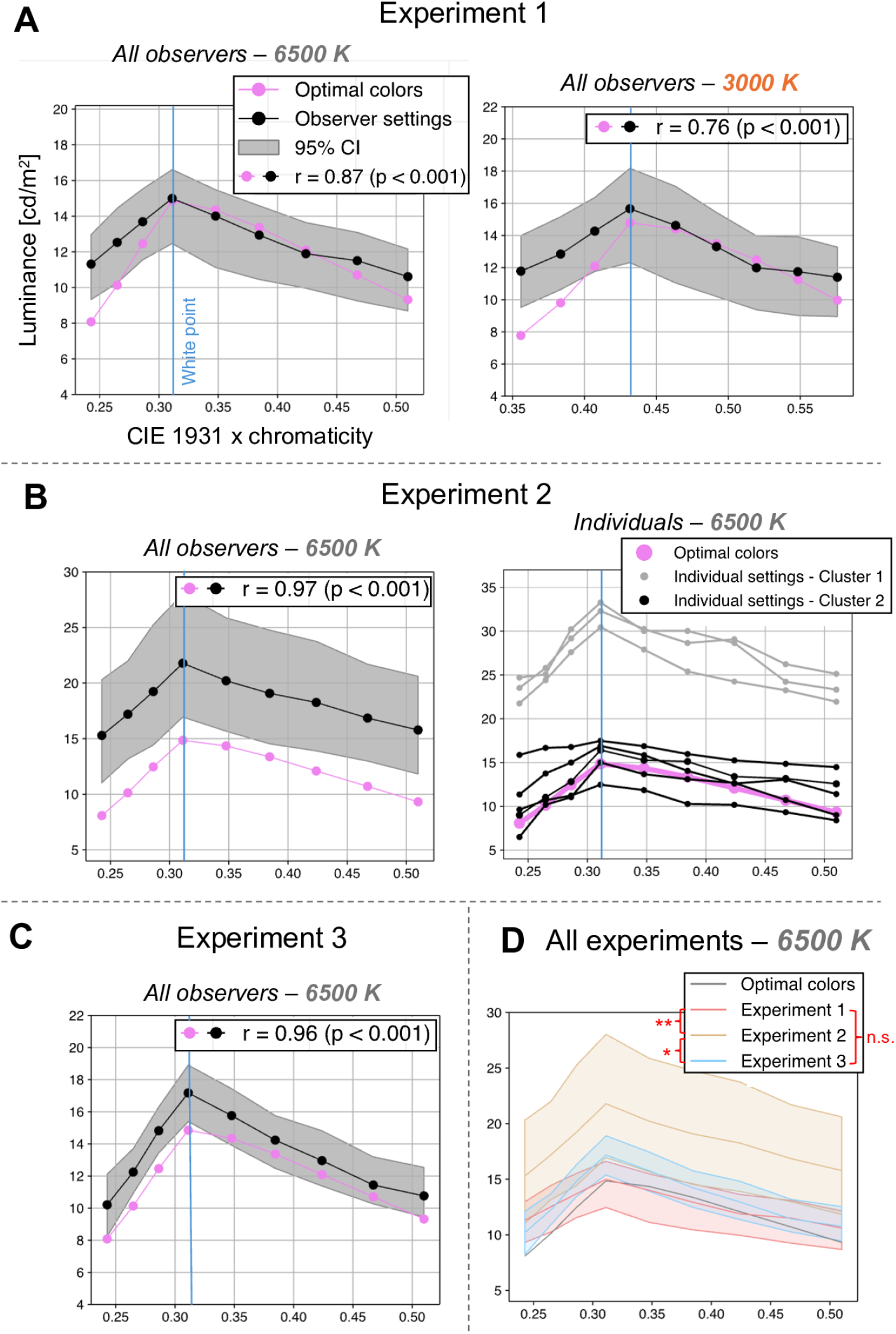
Results of the three experiments. **A**: results of Experiment 1 (averaged across all observers and backgrounds for condition 6500 K on the left and 3000 K on the right). **B**: results of Experiment 2 (averaged across all observers and backgrounds on the left and individual results on the right). **C**: results of Experiment 3 averaged across all observers and backgrounds. **D**: results of the three experiments averaged across all observers and backgrounds plotted on the same chart (only condition with illuminant 6500 K for Experiment 1). Asterisks indicate significant differences between the loci: * *p* < 0.05 and ** *p* < 0.01. Filled colored areas represent 95% confidence intervals. See the Methods section for details on the statistical analysis.

Optimal colors are a well-established concept in the literature and have been presented multiple times, both from theoretical and practical perspectives (e.g., computing optimal color coordinates). Accordingly, we provided the necessary information for understanding the present study, while omitting certain technical details, such as the computational procedures involved in deriving the optimal colors. Instead, we refer the reader to the relevant literature for a more comprehensive overview [2, 13, 16].

### Analysis

All statistical tests, confidence intervals, and *p*-values reported in this study were performed using a two-tailed, non-parametric bias-corrected and accelerated (BCa) bootstrapping method with 10,000 resamples of the eight observers and a significance level of 5% [22]. This approach provides robust estimates that account for potential bias and skewness in the sampling distribution. All *r* -values reported, including those associated with the regression lines in Fig 3A, correspond to Pearson’s correlation coefficients. The statistical significance of these correlations (*p*-values) was assessed using the aforementioned BCa bootstrapping procedure. The regression lines presented in Fig 3A were obtained via standard linear least-squares regression. The accompanying *r* -values represent the Pearson’s correlation coefficients and their significance levels were likewise computed using the aforementioned BCa method.

**Fig 3.**
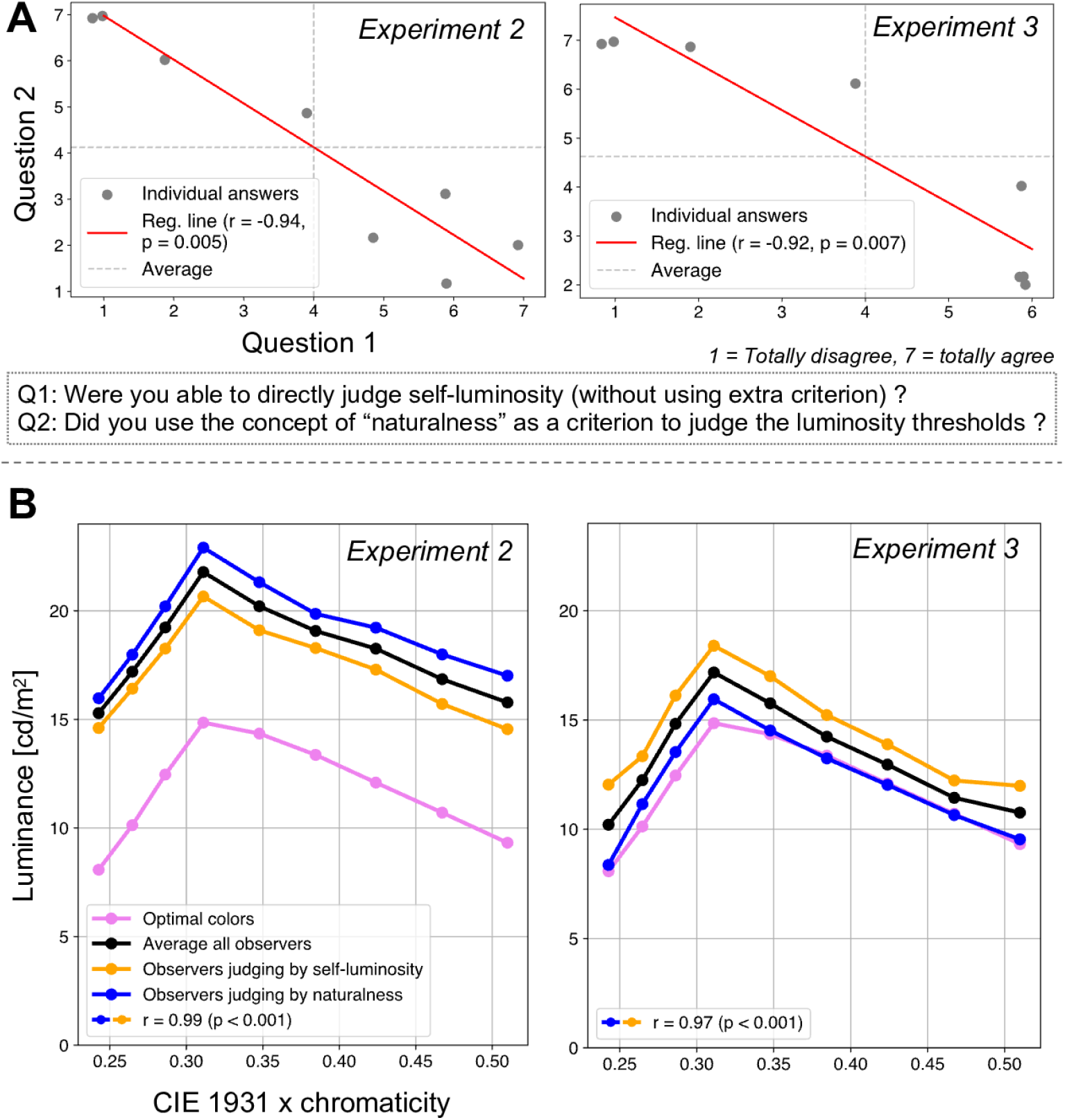
Additional analysis of Experiments 2 and 3. **A**: results of the additional questionnaire for Experiment 2 (on the left) and Experiment 3 (on the right). **B**: comparison and correlation between the settings made by observers who were able to directly judge the luminosity thresholds through the impression of self-luminosity and those who relied on the notion of naturalness to make their judgments (the clustering is based on the responses to the questions presented in panel *A* of this figure) in Experiment 2 (on the left) and Experiment 3 (on the right). See the Methods section for details on the statistical analysis.

## Experiment 1

In Experiment 1, we showed that the optimal colors could predict luminosity thresholds of 2D uniform circles displayed on top of natural backgrounds.

### Stimulus and procedure

Experiment 1 follows the general design presented above. In Experiment 1, observers were tasked with judging the luminosity thresholds of a target presented in different natural backgrounds. The target was a 2D uniform circle with a diameter of approximately 1.29°, overlaid at the center of the background images (see Fig 1C). Observers judged the luminosity thresholds for different chromaticities of the target across all natural backgrounds (see Fig 1A and 1B). To determine the luminosity threshold, they were asked to adjust the luminance of the target using the scroll wheel of a mouse and set it to the point at which the target began to appear self-luminous (see Fig 1E). The luminance was applied uniformly to all pixels of the 2D circular target. The experiment was conducted under two different illuminants: once with stimuli generated under a neutral illuminant with a color temperature of 6500 K and once under an orangish illuminant with a color temperature of 3000 K (see Fig 1B and 1D). Observers completed four sessions under the 6500 K illuminant and four under the 3000 K illuminant, in randomized order. Observers were permitted to take a break between sessions. The experiment was conducted on a single day.

The aim of this experiment was to determine whether the observers’ settings matched the optimal colors predictions. To verify this, we plotted the average settings of all observers across all backgrounds for each chromaticity separately (forming the locus of observers’ settings) along with the optimal colors (forming the locus of optimal colors). We then examined whether there was a correlation between the two loci. We plotted these data on a 2D diagram, selecting the CIE 1931 chromaticity coordinate *x* as the horizontal axis to improve readability of the charts. However, we want to emphasize that the concept of the physical gamut, which is visualized by optimal colors, is inherently 3D, with the CIE 1931 chromaticity coordinates *x* and *y* forming the horizontal plane and luminance representing the elevation (see Fig 1F). We hypothesized a strong correlation between the two loci, supporting the theory of optimal colors predicting luminosity thresholds in these conditions. This expectation was based on prior research demonstrating that the theory holds for uniform circles against abstract backgrounds, provided that the backgrounds allow for the identification of the chromaticity and intensity of the illuminant [2]. Given that natural backgrounds contain rich illumination cues, they were expected to support this effect.

## Results

As anticipated, the results confirmed the validity of the predictions of luminosity thresholds by optimal colors for 2D uniform circles on natural backgrounds. Fig 2A shows the observers’ settings and optimal colors loci separately for the 6500 K and 3000 K illuminant conditions. A strong correlation was found between the two loci (for 6500 K: *r* = 0.87, *p <* 0.001; for 3000 K: *r* = 0.76, *p <* 0.001). Additionally, the shapes of the loci are similar; in other words, the perceived luminosity thresholds match the optimal colors. Consistent with Morimoto et al. [2], the chromaticity of the observers’ peak setting corresponds to the chromaticity of the white point of the illuminant. Indeed, a shift in peak settings was observed between the 6500 K and 3000 K conditions, following the white point of the illuminant. Furthermore, as reported by Morimoto et al. [2], the optimal colors predictions appear to be slightly less precise for the illuminant 3000 K. This discrepancy can be attributed to the theory’s reliance on the estimation of the chromaticity and intensity of the illuminant. Since estimating an orangish illuminant is inherently more challenging and less precise than estimating a neutral illuminant (which corresponds to 6500 K in our case), the reduced accuracy for 3000 K can be expected [2, 15, 16].

## Experiment 2

In Experiment 2, we showed that the optimal colors could predict luminosity thresholds of 3D spheres displayed on top of natural backgrounds.

### Stimulus and procedure

The design, task, and objective of Experiment 2 follow the general design presented above and they were similar to those of Experiment 1, with the primary difference being that the target was a 3D sphere instead of a 2D uniform circle, subtending the same visual angle (see Fig 1C). The aim here was to enhance the naturalness of the stimulus compared to Experiment 1 and to assess whether the optimal colors predicting luminosity thresholds still holds. To create the impression of three-dimensionality, shading was applied to the target from Experiment 1: while the maximum luminance was preserved for a portion of the circle, a circular and directional gradient was introduced to the rest of the surface (see S1 Fig in Supporting information). In the actual experiment, the luminance set by the observer was applied to each pixel of the sphere and multiplied by the shading, such that the area of maximum luminance on the sphere corresponds to the value chosen by the observer, then gradually decreases according to the shading. In Experiment 2, only the 6500 K illuminant was used, and observers completed four sessions. Observers were permitted to take a break between sessions. Experiment 2 was performed on a different day as Experiment 1. An additional modification was introduced in Experiment 2: a questionnaire was administered at the end of the experiment to evaluate the observers’ subjective experience regarding task difficulty and collect spontaneous comments. The questions and details of the questionnaire are presented in Table 1.

**Table 1.**
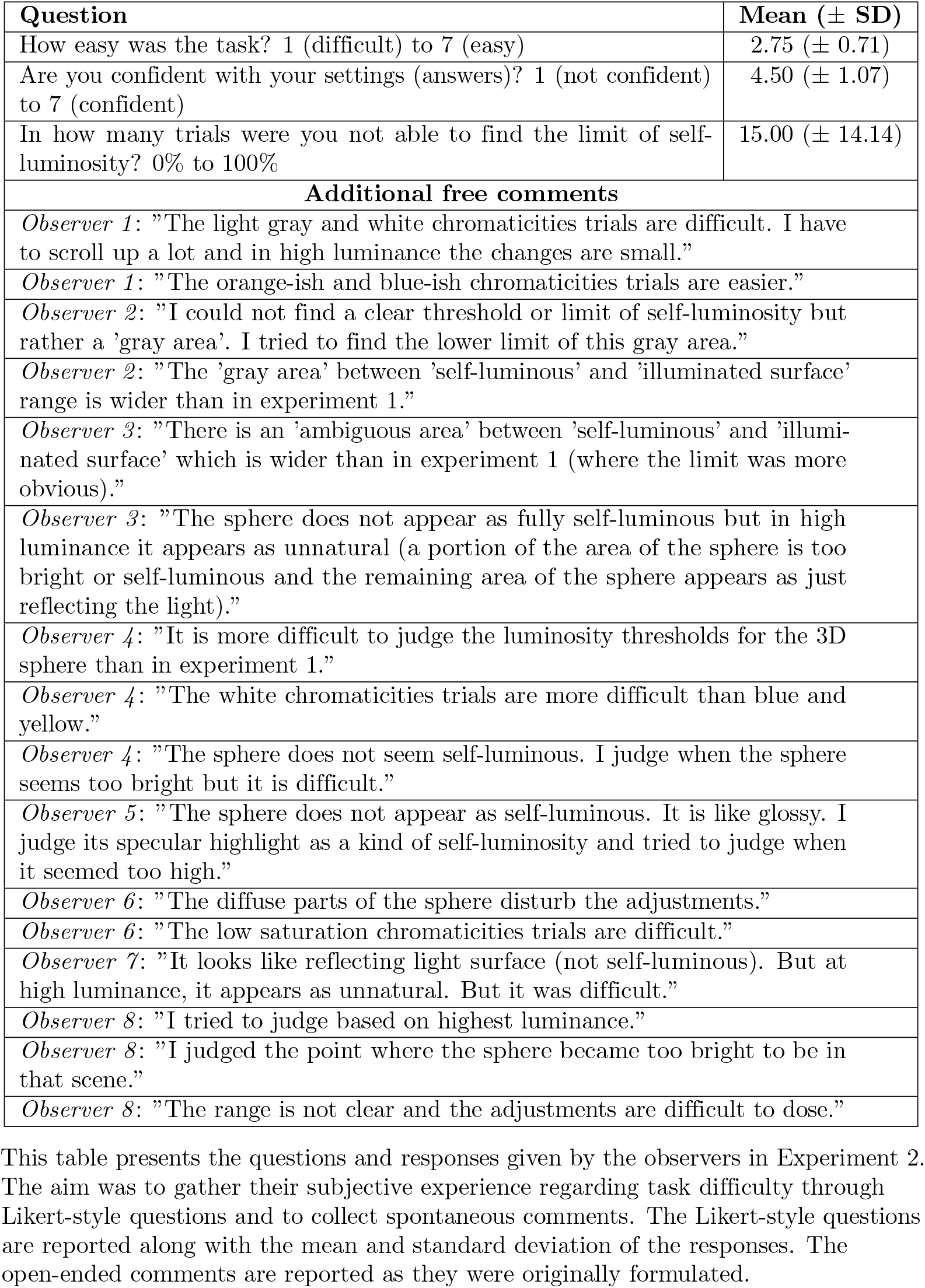
Results of first questionnaire for Experiment 2.

| Question | Mean (± SD) |
| --- | --- |
| How easy was the task? 1 (difficult) to 7 (easy) | 2.75 (± 0.71) |
| Are you confident with your settings (answers)? 1 (not confident) to 7 (confident) | 4.50 (± 1.07) |
| In how many trials were you not able to find the limit of self-luminosity? 0% to 100% | 15.00 (± 14.14) |
| **Additional free comments** |  |
| *Observer 1:* "The light gray and white chromaticities trials are difficult. I have to scroll up a lot and in high luminance the changes are small." |  |
| *Observer 1:* "The orange-ish and blue-ish chromaticities trials are easier." |  |
| *Observer 2:* "I could not find a clear threshold or limit of self-luminosity but rather a 'gray area'. I tried to find the lower limit of this gray area." |  |
| *Observer 2:* "The 'gray area' between 'self-luminous' and 'illuminated surface' range is wider than in experiment 1." |  |
| *Observer 3:* "There is an 'ambiguous area' between 'self-luminous' and 'illuminated surface' which is wider than in experiment 1 (where the limit was more obvious)." |  |
| *Observer 3:* "The sphere does not appear as fully self-luminous but in high luminance it appears as unnatural (a portion of the area of the sphere is too bright or self-luminous and the remaining area of the sphere appears as just reflecting the light)." |  |
| *Observer 4:* "It is more difficult to judge the luminosity thresholds for the 3D sphere than in experiment 1." |  |
| *Observer 4:* "The white chromaticities trials are more difficult than blue and yellow." |  |
| *Observer 4:* "The sphere does not seem self-luminous. I judge when the sphere seems too bright but it is difficult." |  |
| *Observer 5:* "The sphere does not appear as self-luminous. It is like glossy. I judge its specular highlight as a kind of self-luminosity and tried to judge when it seemed too high." |  |
| *Observer 6:* "The diffuse parts of the sphere disturb the adjustments." |  |
| *Observer 6:* "The low saturation chromaticities trials are difficult." |  |
| *Observer 7:* "It looks like reflecting light surface (not self-luminous). But at high luminance, it appears as unnatural. But it was difficult." |  |
| *Observer 8:* "I tried to judge based on highest luminance." |  |
| *Observer 8:* "I judged the point where the sphere became too bright to be in that scene." |  |
| *Observer 8:* "The range is not clear and the adjustments are difficult to dose." |  |

This table presents the questions and responses given by the observers in Experiment 2. The aim was to gather their subjective experience regarding task difficulty through Likert-style questions and to collect spontaneous comments. The Likert-style questions are reported along with the mean and standard deviation of the responses. The open-ended comments are reported as they were originally formulated.

We hypothesized that the theory of optimal colors predicting luminosity thresholds would be validated in Experiment 2, as in Experiment 1. However, based on previously introduced studies, we anticipated that observers’ settings would be relatively higher than in Experiment 1 to compensate for the perceived reduction in luminosity induced by the shading applied to the 3D sphere.

## Results

As expected, the theory of optimal colors predicting luminosity thresholds was confirmed for 3D spheres presented against natural backgrounds. Fig 2B shows the loci of the observers’ settings and the optimal colors, plotted both as an average across all participants and as individual data points. As in Experiment 1, a strong correlation was observed between the optimal colors loci and the averaged observers’ settings (*r* = 0.97, *p* < 0.001), with both exhibiting similar shapes. Moreover, the peak of the observers’ settings corresponds to the white point of the illuminant.

However, the magnitude of the settings is higher in Experiment 2 than in Experiment 1, with an average peak setting of 21.8 compared to 15.0 in Experiment 1. Additionally, the confidence interval is broader in Experiment 2 (range of 11.1) than in Experiment 1 (range of 4.15) for the highest setting, which corresponds to the white point. We assessed intra-observer variability in Experiment 2 and obtained an overall confidence interval with a range of 5.67 (for each observer, confidence intervals were calculated individually for each chromaticity, then averaged to yield a per-observer mean, and finally averaged across all observers to obtain the overall interval). These findings suggest greater inter-observer variability when assessing the 3D spheres. Upon closer examination of individual results (see Fig 2B), two distinct observer groups appeared to emerge. To confirm this, we performed a clustering analysis: the optimal number of clusters was determined using the Elbow method, followed by k-means clustering. This analysis confirmed the existence of two well-separated observer groups, as indicated by the Silhouette Score (0.79) and the Davies-Bouldin Index (0.24). Our interpretation of these clusters suggests that observers in cluster 2 judged the luminosity thresholds based on the highest luminance of the sphere. Their peak setting averages 15.6, closely aligning with the average peak setting from Experiment 1 (15.0). Given that the maximum luminance of the 3D sphere matched that of the 2D uniform circle in Experiment 1, it appears that these observers based their judgments on the brightest portion of the sphere. In contrast, observers in cluster 1 seemed to assess the luminosity thresholds based on the sphere’s average luminance. Their average peak setting is 32, approximately twice those of cluster 2 or Experiment 1. This suggests that these observers compensated for the reduced perceived luminosity caused by shading. Indeed, due to the shading applied to simulate three-dimensionality, the average luminance of the sphere was approximately 50% of the original luminance of the 2D uniform circle in Experiment 1. Thus, these observers appeared to base their judgments on average luminance and adjusted their settings to compensate for the 50% reduction by doubling their original settings. The questionnaire results (see Table 1) support this interpretation. The eight observers in the experiment responded to this questionnaire. Overall, the task was reported as difficult. Three observers specifically mentioned a “gray zone” between the “illuminated surface” and “self-luminous” appearance, and they noted that this gray zone was significantly wider than in Experiment 1. This supports the idea that different perceptual strategies were employed: the ambiguity of this gray zone may reflect whether observers based their judgments on the sphere’s highest luminance or its average luminance. Observers ultimately made a choice in how they interpreted this perceptual boundary.

Additionally, five observers reported that the sphere never appeared self-luminous and three observers reported that they were still able to judge the self-luminosity thresholds based on perceptual naturalness rather than strict self-luminosity. At high luminance levels, the sphere appeared unnatural, as it was perceived as being relatively too bright to be physically plausible in the given scene; this was the breaking point they used for their judgment. Interestingly, a secondary analysis conducted post-experiment and presented in the Discussion section revealed that although these observers did not perceive the stimuli as self-luminous and instead judged them based on “unnatural brightness”, their results were comparable to those of other participants who did perceive self-luminosity (see the Discussion section for further details).

## Experiment 3

In Experiment 3, we showed that the optimal colors could predict luminosity thresholds of in-picture objects in natural backgrounds.

### Stimulus and procedure

The design, task, and objective of Experiment 3 follow the general design presented above and they were similar to those of Experiments 1 and 2. However, the key difference was that the target in this experiment was an “in-picture” element, meaning it was a real-world object already present within the original background rather than a superimposed stimulus (such as the uniform circle in Experiment 1 or the 3D sphere in Experiment 2; see Fig 1C where the targets of Experiment 3 are highlighted in light blue). These target objects were isolated and their chromaticity and luminance were manipulated using masks. Regarding the luminance set by the observer, the luminance adjustment was applied to the masked region corresponding to the target object. To preserve the object’s structure, the adjusted luminance was multiplied by the original pixel luminance values, which had been normalized between 0 and 1 such that the highest pixel value matched the observer’s chosen luminance. These in-picture targets were chosen to represent a wide range of physical properties, including differences in semantic category (e.g., buildings or flowers) and spatial extent (occupying either large or small portions of the image). The aim here was to further enhance the naturalness of the stimulus compared to Experiments 1 and 2 and to evaluate whether the physical gamut theory still holds. In addition, as in Experiment 2, only the neutral illuminant was used. Observers completed four sessions and were permitted to take a break between sessions. Experiment 3 was conducted on the same day as Experiment 2, after a 30-minute break. At the end of the experiment, participants completed the same questionnaire as in Experiment 2 to assess their subjective experience of task difficulty and to gather spontaneous comments. The questions and details of the questionnaire are presented in Table 2.

**Table 2.** Results of first questionnaire for Experiment 3.

| Question | Mean (± SD) |
| --- | --- |
| How easy was the task? 1 (difficult) to 7 (easy) | 5.75 (± 0.71) |
| Are you confident with your settings (answers)? 1 (not confident) to 7 (confident) | 6.25 (± 0.46) |
| In how many trials were you not able to find the limit of self-luminosity? 0% to 100% | 0.00 (± 0.00) |
| **Additional free comments** |  |
| Observer 1: "It is very easy. Small adjustments are more perceptible than in experiment 2." |  |
| Observer 1: "The painted wall background was difficult." |  |
| Observer 2: "The target is part of the scene. It seems natural so it is easy to find the threshold. I can compare with the background. When the luminance is too high, it feels outside the image and unnatural; this is the breaking point that I judged as self-luminous even if it does not look self-luminous." |  |
| Observer 3: "It is much easier than experiment 2 because the 'gray area' range is much narrower." |  |
| Observer 4: "I can identify a top-down effect: I can judge from the shadows like in real life and I know that this is a natural luminance for a real object." |  |
| Observer 4: "It is easy because I can separate the target from the background if the luminance is not matching. It appears as unnatural. This is the strategy that I used." |  |
| Observer 5: "If the object seems self-luminous, it looks like it is outside of the picture (closer to me)." |  |
| Observer 6: "The trials with the image with the small flowers were difficult." |  |
| Observer 6: "The low saturation chromaticities trials were difficult." |  |
| Observer 7: "The trials with the big rose were difficult." |  |
| Observer 7: "Overall, it was easier because the target is inside the image." |  |
| Observer 8: "I judged the self-luminosity as the limit between natural and unnatural. At high luminance, the target seems detached from the background." |  |

This table presents the questions and responses given by the observers in Experiment 3. The aim was to gather their subjective experience regarding task difficulty through Likert-style questions and to collect spontaneous comments. The Likert-style questions are reported along with the mean and standard deviation of the responses. The open-ended comments are reported as they were originally formulated.

Based on previously introduced studies, our hypothesis was that the theory of optimal colors predicting luminosity thresholds would not hold in Experiment 3. We anticipated that, because the in-picture targets were natural objects with textures, they would disrupt the perception of self-luminosity, preventing the same results observed in Experiments 1 and 2 from emerging.

## Results

Contrary to expectations, the results revealed that the theory of optimal colors predicting luminosity thresholds was fully validated even for in-picture targets embedded within natural backgrounds. Fig 2C illustrates the loci of the observers’ settings and the optimal colors, plotted as an average across all participants. As in Experiments 1 and 2, a strong correlation was observed between the optimal colors loci and the observers’ settings (*r* = 0.96, *p* < 0.001), with both exhibiting a similar shape. Furthermore, the peak of the observers’ settings once again corresponded to the white point of the illuminant.

However, in contrast to Experiment 2, the magnitude of the settings in Experiment 3 was not notably higher than in Experiment 1 (average peak setting of 17.1 in Experiment 3 vs. 15.0 in Experiment 1). Likewise, the confidence interval was not notably broader (range of 3.57 in Experiment 3 vs. 4.15 in Experiment 1). This suggests that, unlike in Experiment 2, observers did not adopt different strategies for judging luminosity thresholds (i.e., judging based on the average luminance of the target vs. its maximum luminance). A plausible explanation is that, in Experiment 3, the target was an integral part of the original image, allowing observers to make direct comparisons with the surrounding elements in the scene. The questionnaire results (see Table 2) further support this interpretation. The eight observers in the experiment responded to this questionnaire. The task was reported to be markedly easier than in Experiment 2.

Four observers explicitly stated that the task was straightforward because the target was part of the original scene, enabling them to compare it directly with the background.

Moreover, as in Experiment 2, four observers reported that the target never appeared self-luminous, and three observers reported that they were still able to judge the self-luminosity threshold based on perceptual naturalness. They reported that, at high luminance levels, the target appeared unnatural, as it was perceived as being relatively too bright to be physically plausible in the given scene; this was the breaking point they used for their judgment. Again, the secondary analysis conducted post-experiment and presented in the Discussion section showed that, although these observers did not perceive the stimuli as self-luminous and instead judged them based on “unnatural brightness”, their results were comparable to those of other participants who did perceive self-luminosity (see the Discussion section for further details). In addition, the same observers reported that this unnaturalness was even easier to detect than in Experiment 2. Indeed, when the target’s luminance increased beyond a certain threshold, it appeared to detach from the original image, creating the impression that it was no longer part of its original plane but instead shifted closer to the observer’s eyes. This effect facilitated the identification of the point at which the target became “too bright” or “unnatural”. This phenomenon might also explain why, unlike in Experiment 2, observers did not split into two groups based on different perceptual strategies (i.e., judging based on the average luminance of the target vs. its maximum luminance). Because the target was part of the original image, observers could directly compare it with the background, making judgments about unnaturalness or self-luminosity more straightforward and consistent across participants.

## Discussion

For Experiment 1, our hypothesis was that the optimal colors theory predicting luminosity thresholds would hold in natural backgrounds with targets in the form of 2D uniform circles. This hypothesis was confirmed for two different illuminants, with the theory performing slightly better under neutral light compared to orangish light, reinforcing previous findings under slightly different conditions. For Experiment 2, we hypothesized that the optimal colors predictions would be valid for natural backgrounds with 3D spherical targets, but that observers would set higher thresholds to compensate for shading effects on the sphere. This was only partially confirmed: some observers indeed set thresholds twice as high to compensate for shading over 50% of the sphere’s surface, while others made settings identical to those in Experiment 1. For Experiment 3, we hypothesized that the optimal colors theory predicting luminosity thresholds would not hold, as textured natural objects are never perceived as self-luminous. However, contrary to expectations, this hypothesis was entirely refuted, as the optimal colors could fully predict luminosity thresholds in this scenario, if not more robustly than in Experiment 1. Fig 2D summarizes these results and confirms that the locus of observers’ settings was significantly higher in Experiment 2 than in Experiment 1 (*p* = 0.006) and Experiment 3 (*p* = 0.049). The loci of observers’ settings in Experiment 1 and Experiment 3 did not differ significantly (*p* = 0.37).

The present results provide additional support for the theory of an internal reference in the form of a physical gamut predicting luminosity thresholds, visualized through optimal colors, which was proposed by Morimoto et al. [2] and validated in this study under more natural conditions. Moreover, our findings suggest that luminosity thresholds may encompass both the concepts of self-luminosity and naturalness. Indeed, many observers reported that they could not perceive the targets as self-luminous; instead, they judged them based on a concept of “unnatural brightness”, yet their results were comparable to those of participants who did perceive self-luminosity.

Therefore, when observers are asked to assess the luminosity thresholds of objects that they do not perceive as self-luminous, they appear to rely on the notion of unnaturalness (defined, based on observers’ comments, as “a luminosity level too high for the scene to be plausible”). After analyzing the observers’ settings and responses from the first questionnaire associated with Experiments 2 and 3, we conducted an additional questionnaire to verify this conclusion. The goal of the follow-up questionnaire was to determine whether all observers were unable to directly assess self-luminosity and whether they instead defaulted to a concept of unnaturalness. The details of the questions and results are presented in Fig 3A. The results indicate a strong negative correlation between the ability to directly assess the self-luminosity of targets and the reliance on naturalness as a judgment criterion (for Experiment 2: *r* = -0.94, *p* = 0.005; for Experiment 3: *r* = -0.92, *p* = 0.007). When observers can directly evaluate self-luminosity, they do not rely on the concept of naturalness. However, when they are unable to make this direct assessment, they use naturalness as a reference. The findings suggest that approximately one-third of observers are clearly able to judge self-luminosity and make little or no use of naturalness, another third struggle to assess self-luminosity and rely predominantly on naturalness, while the remaining third fall in between, integrating both aspects into their judgments. Nevertheless, interestingly, the settings aligned with the physical gamut defined by optimal colors for all observers, regardless of how they judged the thresholds. Indeed, we compared and analyzed the observers’ settings by dividing the data into two groups: one group consisted of observers who were able to directly judge the luminosity thresholds based on the impression of self-luminosity, and the other group included those who relied on the notion of naturalness for their judgments. Grouping was determined by the responses to the questionnaire presented in Fig 3A (based on the upper-left and lower-right quadrants defined by the mean of the responses). We then calculated the correlation between the averages of the two groups. Fig 3B illustrates the results of these analyses and confirms a high correlation between the two groups (for Experiment 2: *r* = 0.99, *p <* 0.001; for Experiment 3: *r* = 0.97, *p <* 0.001), suggesting that both strategies lead to similar outcomes. Consequently, this may imply that the concept of the physical gamut similarly envelopes the two dimensions of self-luminosity and naturalness and could be defined as “all physically possible colors in a scene for an object that does not emit light.” Such a gamut may function as an internal reference against which the visual system assesses the plausibility of a color as a surface property; if a color lies outside this range, it may be perceived as self-luminous or unnatural. The two notions of self-luminosity and naturalness appear to be intrinsically linked, differing in semantics (i.e., “an object being a light source” vs. “an object appearing too bright for the scene”) but not in their absolute threshold values, which remain identical and correspond to luminosity thresholds.

Furthermore, whether observers directly judged self-luminosity or relied on unnaturalness, they appeared to use two distinct strategies to assess the luminosity thresholds of more complex natural stimuli (3D spheres): either by considering the target’s average luminance or its maximum luminance. This introduced difficulty, as observers needed to arbitrate between these two strategies, creating ambiguity. However, this difficulty was absent when complex natural objects were directly embedded into the original scene (in-picture targets), as observers could compare them directly to the background. Many observers reported that targets appeared to “come out” of the image when their luminance was too high relative to their surroundings, reinforcing a sense of unnaturalness or self-luminosity and making the task easier by eliminating the need for strategy arbitration (average luminance vs. maximum luminance). This direct comparison to the local surroundings and the effect of the target coming out of the image was not present in Experiment 2, where the 3D sphere, being superimposed onto the background via computer graphics, was already perceived as separate from the image. This distinction might explain why two different strategies were used in Experiment 2 but not in Experiment 3. This observation also raises important questions in other experimental contexts, such as color constancy. For instance, can stimuli involving 3D spheres or 2D uniform circles superimposed on natural backgrounds be effectively used to evaluate color constancy if they are perceived as separate from the original scene and illumination ?

Finally, one might note that the results from Experiment 3 were superior to those from Experiments 1 and 2 in terms of both the correlation between loci and the magnitude of the distance between them. Based on our previous conclusions, we attribute this to the fully natural composition of Experiment 3, which made judgments significantly easier. Whether assessing self-luminosity directly or relying on unnaturalness, observers could use the background and local surroundings for direct comparison. In contrast, in Experiments 1 and 2, targets superimposed via computer graphics introduced ambiguity and increased task difficulty, as the target was not necessarily perceived as an integral part of the background. Similarly, Morimoto et al. [2] obtained results superior to those of our Experiment 1, but their stimuli consisted entirely of 2D uniform circle patterns (both background and target), allowing targets to be perceived as embedded within the background due to their uniform depth and planar alignment. This supports the notion that mixed stimuli, combining real backgrounds with computer-generated targets, introduce ambiguity in judgments; a factor absent in fully natural or fully abstract stimuli, leading to a task easier to conduct in these conditions and superior results. These results, along with the observations described in the previous paragraph, suggest that the consistency and naturalness of an image modulate the way a stimulus is perceived and may influence task performance in psychophysical experiments.

Our study has several limitations that future research could address. First, due to our intention to closely replicate the experimental conditions of Morimoto et al. [2], we used a limited set of backgrounds, illuminations, and target chromaticities. Morimoto et al. [2] demonstrated that the physical gamuts are less accurately predictive when target chromaticities or illuminations are highly unnatural, such as green or magenta. Second, due to the already extensive duration of the experiments, we tested only one illumination color in Experiments 2 and 3. This choice was guided by findings from both Morimoto et al. [2] and our own Experiment 1, which already support extensively the validity of the physical gamut theory under different illumination conditions and suggest that observers adapt the gamuts to the illumination color. We considered unnecessary to re-demonstrate it in Experiments 2 and 3. Despite these first two limitations, the focus of the present study was on evaluating the optimal colors predictions using more naturalistic stimuli while closely adhering to the design of Morimoto et al. [2] in order to minimize the number of secondary factors that could lead to differing results; we determined that the combinations of target chromaticities and illuminations were sufficient to test our hypotheses. Nonetheless, future studies could verify whether the optimal colors predictions hold across a greater diversity of backgrounds, target chromaticities, and illuminant colors, as this remains unclear. Third, we have focused solely on the optimal colors theory and the concept of the physical gamut to explain and define the luminosity thresholds. We took this stance because it is the most recent theory, and it has been shown to be the most effective and superior to other theories [2]. However, future studies could explore other previous theories that explained luminosity thresholds based on the literature previously cited. Finally, we tested our theory with only eight observers. It would be valuable to assess these findings in a larger sample, particularly to further validate the different judgment strategies proposed in this study (i.e., average vs. maximum luminance of the target and self-luminosity vs. unnaturalness).

Another point of discussion concerns the process by which the physical gamut is internalized. As proposed by Morimoto et al. [2] and adopted in the present study, this internal reference is thought to be constructed by the human visual system through empirical observation of the real world and the surface colors encountered in everyday life. However, this theory does not address whether this construction occurred over generations through biological evolution and historical population movements, or whether it is sensitive to changes on a more recent timescale (a few hundred years), or even to the exposure experienced over the course of a single human lifetime. Potentially, these gamuts and the results of these luminosity thresholds judgements could differ significantly across populations from different regions of the world, depending on the statistical distribution of colors to which they are exposed. Future studies could consider exploring this direction as well. Nevertheless, in a slighlty different but similar matter, it has been shown that such differences do not exist with regard to color discrimination or color preferences, even between regions of the world that differ greatly in terms of environment and color distribution [23].

In conclusion, our findings lend additional support to the theory that luminosity thresholds are guided by an internal reference, conceptualized as a physical gamut that the human visual system builds by empirical observations of the real-world and which is visualized through optimal colors as proposed by Morimoto et al. [2], and demonstrate its validity under more naturalistic conditions. Our results suggest also that luminosity thresholds may integrate both self-luminosity and naturalness notions. Consequently, the concept of the physical gamut may also encompass these two dimensions and be defined as “all physically possible colors in a scene for an object that does not emit light.” This gamut may serve as an internal reference used by the visual system to judge whether a color is plausible as a surface property; when a color falls outside this range, it is more likely to be perceived as self-luminous or unnatural. These findings can have profound potential implications for both applied fields (e.g., AR, projection mapping) and fundamental science (i.e., understanding human visual mechanisms).

## Acknowledgments

This work was supported by JST SPRING Grant Number JPMJSP2180 to KD, and JSPS KAKENHI Grant Number 23K28174 and 21KK0203 to TN.

## Supporting information

**Fig S1.**
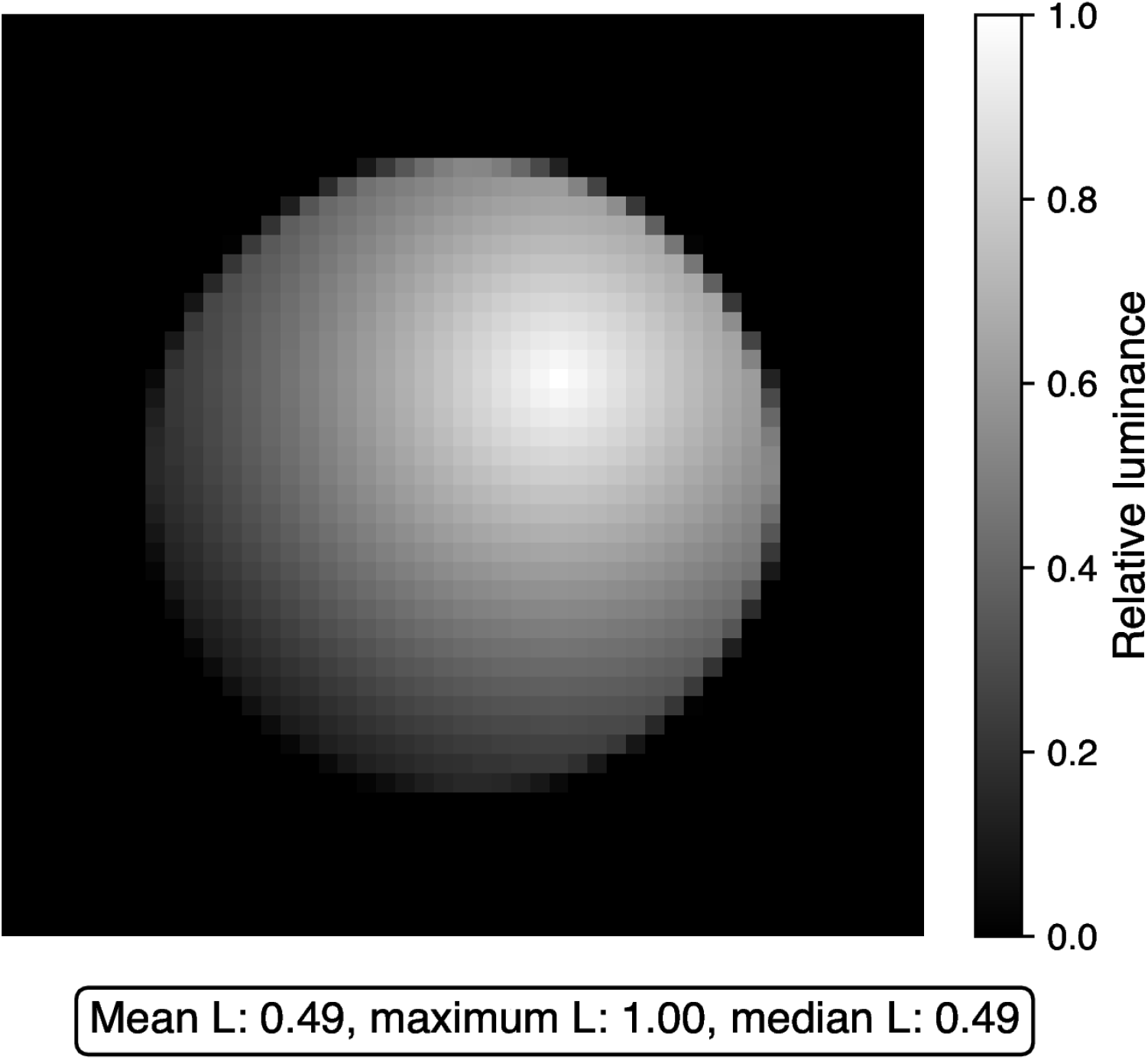
Shading of target in Experiment 2. This represents the shading applied to the circle to create an impression of 3D sphere in Experiment 2. The shading effect is applied to the original circle by scaling each pixel’s luminance based on its distance *d* in pixels from the highest luminance point at (*c*_*x*_ + 5, *c*_*y*_ − 5), where (*c*_*x*_, *c*_*y*_) is the center of the circle in pixels, using the formula: shaded = original × (1 − *d/*(*r* · *s*)), where *r* = 16 is the radius in pixels and *s* = 1.6 is a scalar value representing the shadow strength. The gradient is presented in the form of relative luminance, which was then multiplied by the luminance set by the observer during the completion of the experimental task, in order to create the desired final shaded sphere at different luminance intensities based on observer’s adjustments.

## References

1. Katz D. The world of colour. Routledge; 2013.

2. Morimoto T, Numata A, Fukuda K, Uchikawa K. Luminosity thresholds of colored surfaces are determined by their upper-limit luminances empirically internalized in the visual system. Journal of vision. 2021;21(13):3–3.

3. Evans RM. Fluorescence and gray content of surface colors. Journal of the Optical Society of America. 1959;49(11):1049–1059.

4. Evans RM, Swenholt BK. Chromatic strength of colors: dominant wavelength and purity. Journal of the Optical Society of America. 1967;57(11):1319–1324.

5. Evans RM, Swenholt BK. Chromatic Strengths of Colors, Part II. The Munsell System. Journal of the Optical Society of America. 1968;58(4):580–584.

6. Evans RM, Swenholt BK. Chromatic strength of colors, III. Chromatic surrounds and discussion. Journal of the Optical Society of America. 1969;59(5):628–634.

7. UlIman S. On visual detection of light sources. Biol cybernetics. 1976;21:205–212.

8. Bonato F, Gilchrist AL. The perception of luminosity on different backgrounds and in different illuminations. Perception. 1994;23(9):991–1006.

9. Speigle JM, Brainard DH. Luminosity thresholds: effects of test chromaticity and ambient illumination. Journal of the Optical Society of America A. 1996;13(3):436–451.

10. Uchikawa K, Koida K, Meguro T, Yamauchi Y, Kuriki I. Brightness, not luminance, determines transition from the surface-color to the aperture-color mode for colored lights. Journal of the Optical Society of America A. 2001;18(4):737–746.

11. MacAdam DL. Maximum visual efficiency of colored materials. Journal of the Optical Society of America. 1935;25(11):361–367.

12. MacAdam DL. The theory of the maximum visual efficiency of colored materials. Journal of the Optical Society of America. 1935;25(8):249–252.

13. Uchikawa K, Fukuda K, Kitazawa Y, MacLeod DI. Estimating illuminant color based on luminance balance of surfaces. Journal of the Optical Society of America A. 2012;29(2):A133–A143.

14. Fukuda K, Uchikawa K. Color constancy in a scene with bright colors that do not have a fully natural surface appearance. Journal of the Optical Society of America A. 2014;31(4):A239–A246.

15. Morimoto T, Fukuda K, Uchikawa K. Effects of surrounding stimulus properties on color constancy based on luminance balance. Journal of the Optical Society of America A. 2016;33(3):A214–A227.

16. Morimoto T, Kusuyama T, Fukuda K, Uchikawa K. Human color constancy based on the geometry of color distributions. Journal of Vision. 2021;21(3):7–7.

17. Kuriki I. Effect of material perception on mode of color appearance. Journal of vision. 2015;15(8):4–4.

18. Kuriki I, Uchikawa K. Limitations of surface-color and apparent-color constancy. Journal of the Optical Society of America A. 1996;13(8):1622–1636.

19. Wyszecki G, Stiles WS. Color science: concepts and methods, quantitative data and formulae. John wiley & sons; 2000.

20. Foster DH, Reeves A. Colour constancy failures expected in colourful environments. Proceedings of the Royal Society B. 2022;289(1967):20212483.

21. Foster D, Amano K, Nascimento S. Fifty hyperspectral reflectance images of outdoor scenes; 2022.

22. Efron B. Better bootstrap confidence intervals. Journal of the American statistical Association. 1987;82(397):171–185.

23. Skelton AE, Maule J, Floyd S, Wozniak B, Majid A, Bosten JM, et al. Effects of visual diet on colour discrimination and preference. Proceedings of the Royal Society B. 2024;291(2031):20240909.

